# The strength of a gene’s first 5’ splice site is a key determinant of its expression

**DOI:** 10.64898/2026.09.28.754465

**Authors:** Yuta Sakai, Michael P. McGurk, Emma J. Kowal, Elaine Huang, Christopher B. Burge

## Abstract

The control of gene expression is central to cellular physiology and to various biotechnological applications. In higher eukaryotes, synthesis of most messenger RNAs involves removal of introns during splicing. Intron-containing genes are often more highly expressed than intronless versions, despite generating the same mRNA transcripts, suggesting that the splicing of a gene’s transcripts positively impacts the gene’s expression. Termed intron-mediated enhancement (IME), this phenomenon has been observed in animals, plants, and fungi, with enhancement ranging from 2- to 100-fold and the strongest effects observed for promoter-proximal introns^1–3^. Here, we assayed thousands of natural human introns, covering all distinct human 5’ splice site (5SS) motifs and thousands of 3’ splice sites (3SS), for their ability to enhance (or repress) expression of a reporter gene. The results confirmed previous observations that intronic UUUU motifs augment the magnitude of IME, and revealed two fundamental features of IME: (i) only introns spliced by the major spliceosome elicit IME, with minor introns often repressing expression; and (ii) the magnitude of IME increases with the strength of the 5SS motif, but is independent of 3SS strength. Our findings imply that genetic variants that alter the strength of the first 5SS of genes may alter their expression by modulating IME without necessarily impacting splicing. We confirm this expectation using human population genetic data and further show that disease-relevant perturbations of U1 snRNP components that enhance recognition of subsets of first 5SS up-regulate expression of the associated genes. These findings demonstrate that recognition of the first 5SS is vitally important to the expression of human genes.

## INTRODUCTION

RNA splicing often occurs co-transcriptionally, with the spliceosome assembling on the nascent RNA as it exits RNA polymerase II, providing opportunities for the splicing and transcription machineries to interact and potentially influence one another. Spliceosomal components may promote gene expression through interactions with general transcription factors at the promoter^4^, by facilitating release of paused polymerase^5^, enhancing elongation rates in introns^6^, or inhibiting recognition of intronic termination signals^7^. The exon-junction complex (EJC) deposited during the splicing reaction further promotes packaging, export, and translation of the mRNA, sometimes influencing transcript stability^8^. These effects, and perhaps others, have led to the frequent observation that presence of an intron boosts gene expression, a phenomenon termed intron-mediated enhancement (IME).

But what principles govern whether and how strongly an intron will exert IME? Work in a variety of organisms suggests that the effect of an intron increases with its proximity to the promoter^2,3,9^. Quantitative studies of human cells have supported that the predominant effect of introns on gene expression is by increasing mRNA production rather than effects on mRNA stability or nucleocytoplasmic export^10^. Across mammals, the evolutionary emergence of new exons inside 5’ untranslated region (UTR) introns can trigger the emergence of nearby, upstream transcription start sites, whose activity was found to be dependent on the splicing of the new exon^11^. While positive effects on transcription and more modest impacts on export have been observed in many genes^9,12^, other mechanisms can contribute in some cases, e.g., effects on 3’ end processing at a weak polyadenylation signal in the human beta-globin gene^13,14^. To investigate the sequence determinants for IME, we previously performed a massively parallel reporter assay (MPRA), interrogating the effects on expression of 19,000 randomized intronic sequences flanked by fixed strong splice sites^15^. That study found that nearly all random introns enhanced expression relative to an intronless control and that presence of U-rich motifs in introns was associated with stronger IME. However, natural intron sequences are far from random, having been shaped by selection acting on splicing speed and fidelity and perhaps effects on expression. Another wrinkle is that two different spliceosomes coexist in most eukaryotes, with the ‘major’ spliceosome responsible for most introns, and a compositionally distinct ‘minor’ spliceosome responsible for splicing of a small minority (∼0.3%) of human introns with distinct splice site motifs^16^.

With these considerations in mind, we designed a new internally controlled MPRA to assess IME across diverse natural human introns, with over 2,000 distinct intron bodies and over 3,000 distinct human splice site pairs, encompassing all human 5’ splice site (5SS) motifs and diverse 3’ splice site (3SS) motifs. The results revealed fundamental features of IME, including a stark distinction between the major and minor spliceosomes, strong dependence on 5SS strength but not 3SS strength, and implicate the strength of a gene’s first 5SS of a gene as a key determinant of its expression.

## RESULTS

### A natural intron screen yields robust measurements of IME for individual introns

To investigate the effects of natural introns on gene expression (Fig. 1a), we designed a diverse library of intron sequences from the human genome (Fig. 1b). We employed a two-step cloning strategy to generate a plasmid library from array-synthesized DNA oligonucleotides pools encoding intron sequences and associated barcodes (Supplementary Fig. 1a). This approach yielded intron-containing EGFP and intronless dTomato (dTom) reporter genes with identical 12-nt barcodes in their 5’ UTRs (Fig. 1b). This design enables comparison of the expression of the intron-containing EGFP to that of the intronless dTom expressed from the same plasmid. Both genes have matching UbC promoters, 3’ UTRs and polyadenylation signals. To control for any baseline expression/amplification differences between EGFP and dTom, we also included plasmids containing dTom paired with an intronless EGFP (plasmids and primers used are listed in Supplementary Tables S1-S3).

**Figure 1:**
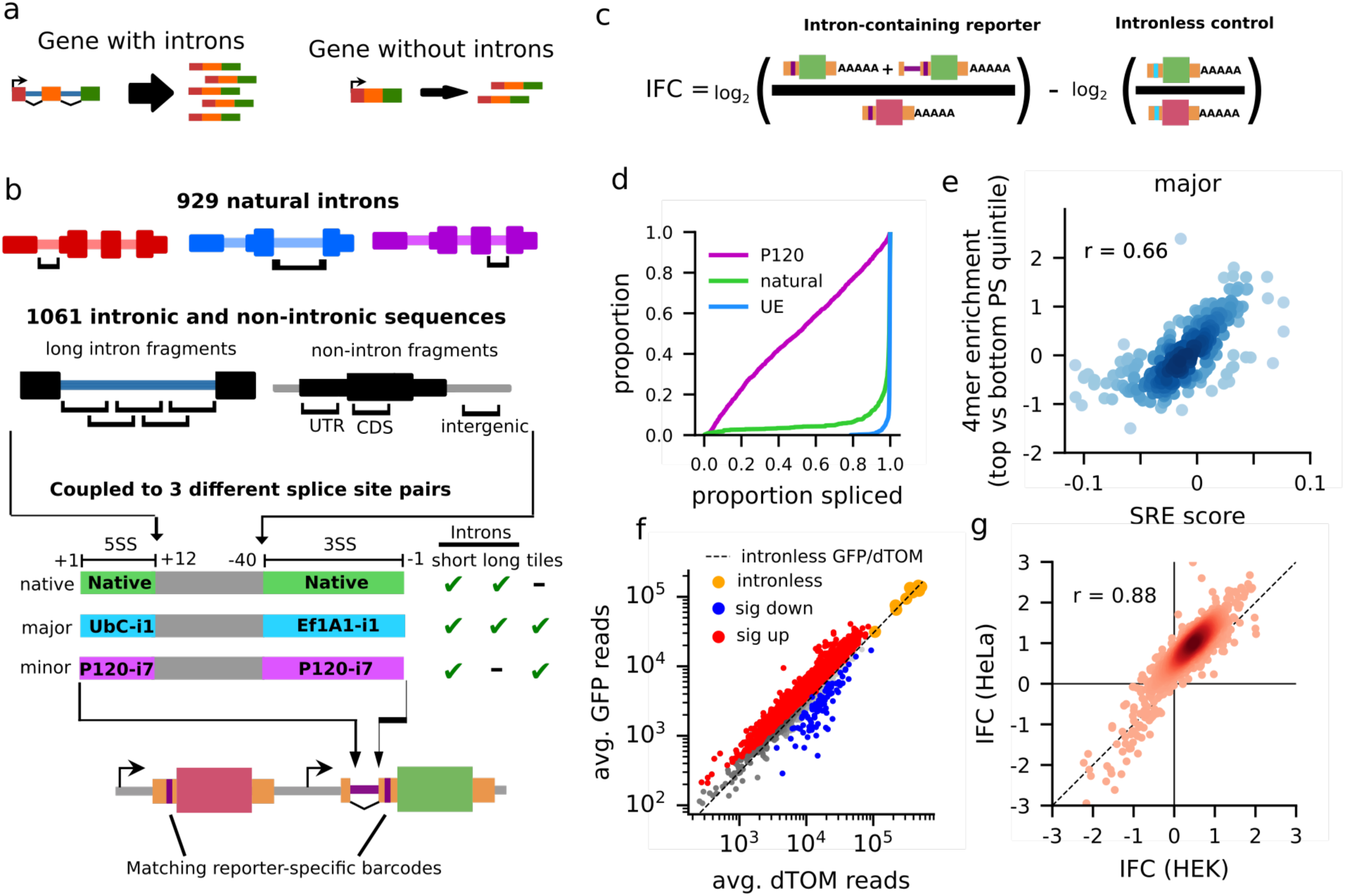
Design of a natural intron library for analysis of IME. **a)** Illustration of intron-mediated enhancement. **b)** Overview of the natural intron-library. **c)** Illustration of how IFC is computed by comparing the EGFP/dTom ratio of intron-containing reporters to that of intronless reporters. **d)** The proportion spliced of natural introns with their native splice sites, strong major splice sites (UE), and strong minor splice sites (P120). **e)** The association between the average SMsplice intronic SRE score of tetramers and their log2-enrichment in the most included introns relative to the bottom 20% included introns (top versus bottom quintiles) for introns with native splice sites. **f)** The EGFP and dTom reads for each construct in HEK, averaged across replicated. The dashed diagonal line indicates the mean EGFP/dTom ratio in the intronless controls; introns that significantly increase or decrease EGFP expression relative to these controls are denoted in red and blue, respectively. **g)** Comparison of IFC measured in the HEK and HeLa transfections.

We selected 847 short introns (≤190 nt) which could be synthesized in their entirety as oligo pools, chosen to span a wide range of base compositions from genes well expressed in HEK293 cells (Supplementary Fig. 1b,c) and intron position of origin, as well as 82 additional long introns which were individually synthesized. We further included 660 genomic fragments, including segments tiling across 17 long introns, as well as 401 non-intronic sequences extracted from untranslated regions (5′ and 3′ UTRs), coding sequences (CDS exons), and intergenic regions. We made single-nucleotide substitutions (preserving base composition) to any sequences containing restriction sites used for cloning or potential start codons (ATG) to enable cloning and to limit biases related to translation and nonsense-mediated mRNA decay (NMD), respectively. Thus, our design minimizes the potential for NMD by restricting splicing to the 5’ UTR, eliminating the potential for alternate translation start sites/reading frames, and retaining a constant short 3’ UTR. It also minimizes the potential for effects of splicing on 3’ end processing by placement of the intron near the 5’ end of the gene and use of the strong UbC polyadenylation signal.

To assess the contributions of splice sites to IME, all natural short introns were synthesized with three pairs of splice sites: native, strong major (*UbC* i1 5SS + *EF1A* i1 3SS, referred to as “UE”), and strong AT-AC minor splice sites (*NOP2/*P120 i6). Each short intron/splice-site combination was coupled to three distinct barcode sequences, to increase robustness and control for potential barcode effects. We performed long-read sequencing (Oxford Nanopore) on the resulting plasmid library to assess its quality and confirm that the paired dTom and EGFP barcodes matched as expected. Only constructs with the intended intron sequence and expected matched barcodes were included in the downstream analyses.

To assess IME, we transfected this library into both HEK293 and HeLa cells with two independent transfection replicates each, using transient transfection to maintain library diversity. Amplicon libraries were prepared from the 5′ UTR region and deep sequenced to quantify the three mRNA species expressed from each barcoded construct: dTom mRNA, spliced EGFP mRNA, and unspliced EGFP mRNA. We assessed the proportion spliced (PS) for each intron from the spliced and unspliced EGFP reads. To assess the effect of an intron on expression, we compared the mRNA expressed from the intron-containing EGFP versus that of barcode-matched intronless dTom and normalized this EGFP/dTom ratio to the corresponding ratio from intronless control plasmids (Fig. 1c). We define the log_2_ of this ratio as the “intron-associated fold-change” (IFC), which assesses the impact (positive or negative) of the presence of an intron on gene expression, controlling for any differences in expression, amplification, library prep or sequencing between the EGFP and dTom genes. Thus, positive IFC values represent IME.

Consistent with our previous findings, nearly all introns with the strong major UE splice sites were efficiently spliced. Introns with their native splice sites were often well-spliced, but displayed greater variation in splicing – while use of minor splice sites from the P120 intron often decreased splicing efficiency (Fig. 1d, Supplementary Fig. 1d) with a modest level (1.6%) of cryptic splicing at major splice sites observed (Supplementary Fig. 1e, f). For major introns with native splice sites, the differences in sequence composition between well-spliced and poorly-spliced introns were highly consistent with previously identified splicing regulatory elements^17^ (Pearson’s *r*=0.66, Fig. 1e).

To investigate the determinants of IME and exclude any confounding effects of incomplete splicing, we focused on well-spliced introns. Defining “percent spliced” (PS) as the ratio of the number of spliced reads to the total of all amplicon reads (spliced + unspliced), we restricted further analyses to introns with PS > 0.99. Most well-spliced natural introns displayed robust and reproducible IME, with 85% and 95% of the intron-containing EGFP constructs being more highly expressed than intronless controls (at 10% FDR) in HEK and HeLa, respectively (Fig. 1f, Supplementary Fig. 1g,h). The IFC values of identical introns were strongly correlated across two independent transfections of each cell type in HEK (Pearson’s *r* = 0.97), and in HeLa (Pearson’s *r* = 0.94, Supplementary Fig. 1i). IFC values were also well correlated between HEK and HeLa cells (Pearson’s *r* = 0.88), with generally stronger IME observed in HeLa than in HEK cells (Fig. 1g). The magnitude of enhancement observed in this system – rarely exceeding 4-fold – was more modest than that observed in our previous, random-intron screen, where 8-fold enhancement was common^15^. This difference may reflect changes made to the plasmid design to reduce cryptic splicing potential, minimize library dropout during transfection, and more completely match the UTR sequences of the dTom and EGFP reporters. Sequences, PS and IFC values, etc., for individual introns are listed in Supplementary Table S4.

### Intron composition and U-rich motifs impact IME

Having confirmed the robustness and reproducibility of our screen, we asked which intronic features drive intron-specific IME. Intron bodies (i.e. sequences of introns excluding their splice sites) showed consistent IFC (Pearson’s *r*=0.57) when coupled to UE or native splice sites (Fig. 2a), indicating that the composition of the intron body affects the strength of IME independent of the splice sites^1,15^. However, replacing an intron’s native splice sites with the UE strong major splice sites typically resulted in increased IFC (Fig. 2b, Supplementary Fig. 2a), suggesting additional contributions of the splice sites (explored below).

**Figure 2:**
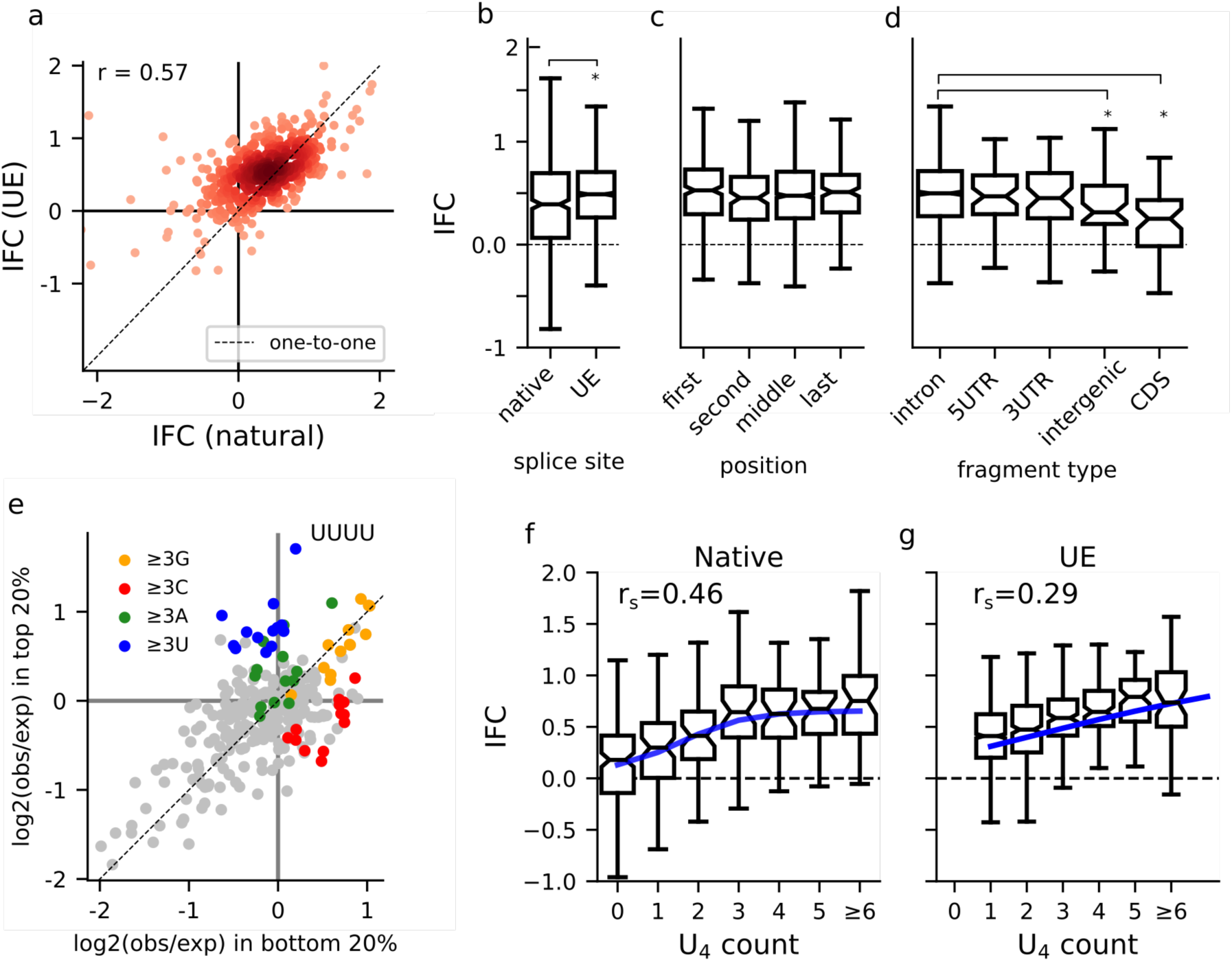
Intron features associated with IME. **a)** The IFC of natural introns with their native splice sites and with those splice sites replaced with strong major (UE) splice sites. **b,c,d)** Box plots of the IFC distributions for intron bodies in the library broken down by native or UE splice sites (**b**), ordering within the gene (**c**), annotated region of origin (**d**): asterisks indicate that the bootstrapped 94%-CI for the difference in means does not overlap zero (10,000 resamples). **e)** The enrichment of tetramers in introns from the top quintile of IFC compared to the bottom quintile. **f, g)** The association between IFC and the number of U_4_ tetramers in the intron’s sequence for introns with native (**f**) and strong major splice sites (**g**); the association between U_4_ count and IFC is summarized with Spearman’s rank correlation (r_s_) and the trendline is a sigmoid with the steepness, midpoint, and asymptotes fit to the observations by minimizing the squared error.

Several previous studies found that introns near transcription start sites (TSS) – typically first introns – elicit stronger IME^1–3^. To ask whether first introns are enriched for features that promote IME, we assessed IME activity for introns originating from different positions in their native gene (first, second, middle, or last). No strong effects of intron position of origin were observed (Fig. 2c, Supplementary Fig. 2a), suggesting that it is the location of first introns rather than some unique sequence property that contributes to IME. Similarly, introns originating from UTRs, from coding regions of genes, or from non-coding RNA genes (Supplementary Fig. 2b), as well as non-intronic fragments derived from 5’/3’-UTR or intergenic regions, all elicited comparable IME to first introns (Fig. 2d, Supplementary Fig. 2a). However, use of coding sequence fragments as intron bodies elicited weaker IME, potentially reflecting differences in sequence composition shaped by coding constraints (Fig. 2d, Supplementary Fig. 2a). Together these observations indicate that intron bodies derived from diverse non-coding sequences are compatible with IME, but that intron sequence composition can influence the magnitude of IME.

We next asked what sequence features might underlie these intron body contributions to IME. Examining differences in tetramer composition between introns in the top and bottom quintiles of IFC, we noted that high-IFC intron bodies were enriched for U-rich tetramers (sequences containing ≥ 3 Us), with UUUU (“U_4_”) being the most enriched tetramer (Fig. 2e, Supplementary Fig. 2c), consistent with our previous findings^15^. These effects of U-rich motifs are general to intron bodies rather than the polypyrimidine tract (PPT) per se, as they do not show positional enrichment near the 3SS (Supplementary Fig. 2d, e). Furthermore, their association with IME was still observed when the native 3SS (including PPT) were replaced with a fixed strong major 3SS (Fig. 2f,g, Supplementary Fig. 2f,g). Each additional U_4_ motif in a natural intron body – up to about three or four motifs – was associated with a +0.15 increase in IFC (bootstrapped 94% CI: +0.11 to +0.19). The per-U_4_ effect on IFC was more modest when the native splice sites were replaced with strong UE splice sites. In general, there appears to be a sub-additive relationship between the effects of splice site strength, favorable cellular context and U_4_ motifs on IME in the sense that each of these impacts IME more strongly in the absence of the others. Specifically, introns lacking U_4_ motifs showed lower IFC with their native splice sites than with UE splice sites, but introns with three or more U_4_ motifs exhibited comparable IFC regardless of the splice sites (Fig. 2f, g). Similarly, the number of U_4_ motifs impacted IME less in HeLa cells (Supplementary Fig. 2f, g), where IME was higher overall.

### Strong 5’ splice sites promote IME

The observation that replacing an intron’s native splice sites with strong UE splice sites leads to increased IME suggests that the splice sites themselves may drive IME. Soon after their emergence from the RNA PolII exit tunnel, the 5SS and PPT/3SS motifs (Fig. 3a) typically recruit U1 snRNP and the U2AF heterodimer, respectively^18^. These roles in the early stages of intron recognition may position these motifs to influence IME. We therefore examined how IME varied among introns with their native splice site motifs to explore the role of splice site strength. Among natural introns with their native splice sites, 508 were well-spliced with PS > 0.99. Estimating the strengths of these splice sites using MaxEntScan^17,19^, most had strong (>8 bits) or moderate (4 to 8 bits) SS, with few weak (<4 bits) SS, as expected for well-spliced introns. This set encompassed more 3SS diversity than 5SS diversity, both because 3SS motifs are longer and more variable, and because the last three exonic bases and the first two intronic bases of the 5SS had fixed consensus bases (CAG/GT) in our library (Supplementary Fig. 3a). Some 88 unique 5SS motifs were represented in these well-spliced introns, varying across the +3 to +6 positions, with strong motifs typically occurring in multiple introns and moderate motifs being represented in one or two introns each (Supplementary Fig. 3b).

**Figure 3:**
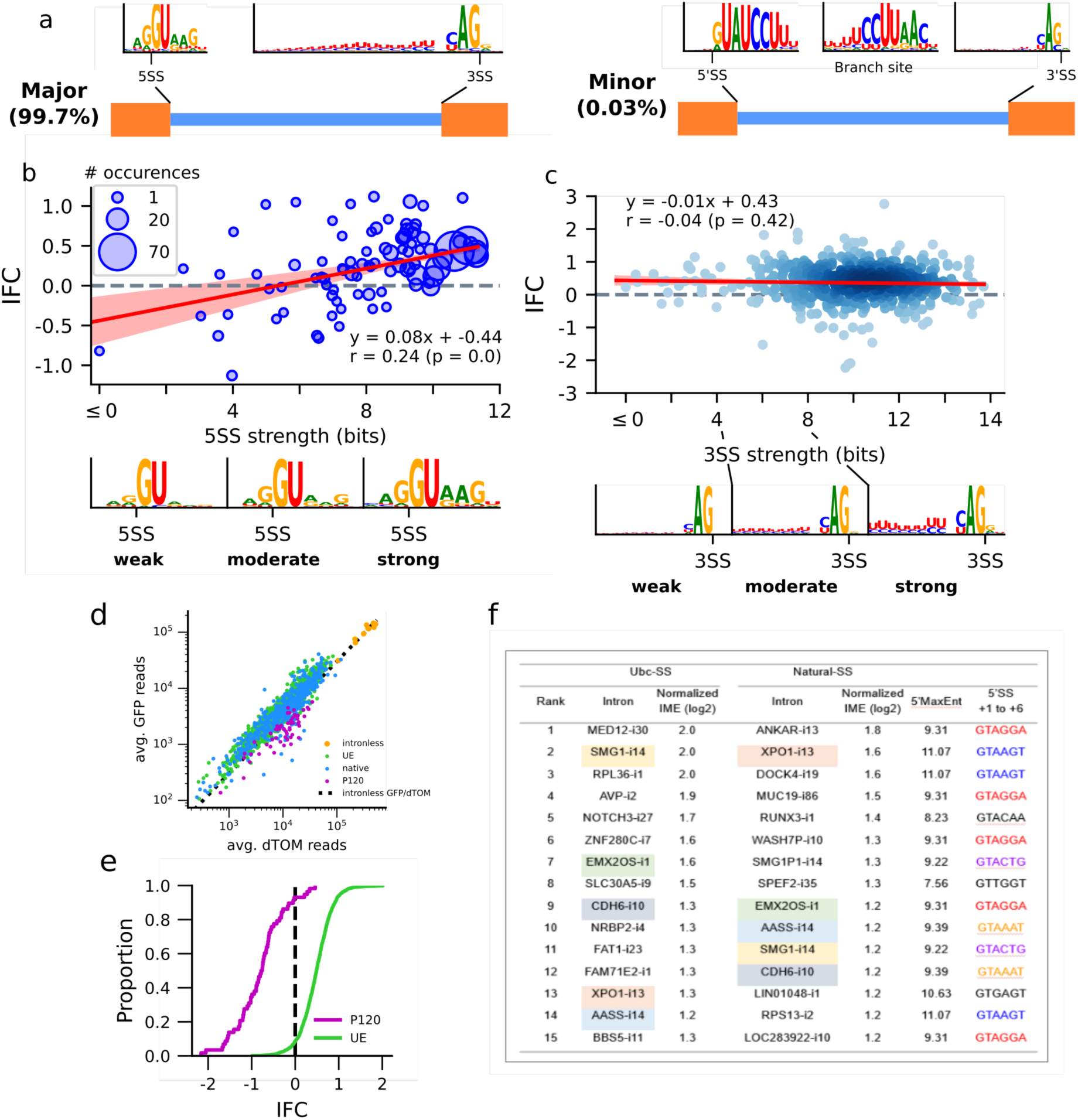
Splice site features associated with IME. **a)** The key signal sequences and their motifs in major (top) and minor introns (bottom). **b, c)** Top: The association between 5SS (**b**) and 3SS (**c**) strength and IME, with IFC averaged across all introns with a given 5SS motif in the library; the size of the dots indicates the number of occurrences. Trend lines are linear regression fits, with 94%-intervals constructed by bootstrapping splice sites 10,000 times. Bottom: sequence logos of weak (< 4 bit), moderate (4-8 bit) and strong (>8 bit) 5SS (**b**) and 3SS motifs (**c**) from the human genome for reference. The 3SS motifs are truncated for compactness (see supp xx for full motifs). **d)** The average EGFP and dTom reads for each construct in HEK, colored by splice site type. The dashed diagonal line indicates the mean GFP/dTom ratio in the intronless controls. **e**) The cumulative distributions of IFC for introns with strong major splice sites (UE) and with strong minor splice sites (P120). **f**) The top 15 enhancing introns identified by the library, with IFC values. Intron names are colored to show matches between UbC-SS and Natural-SS columns. 5’SS motifs (+1 to +6 positions only as –3 to –1 are universally CAG) are colored to show identical motifs.

Natural introns with strong 5SS motifs consistently elicited IME comparable to that observed with the strong *UbC* 5SS, while moderate 5SS motifs were associated with lower levels of IME (Fig. 3b, Supplementary Fig. 3c). These effects on gene expression appeared to increase with splice site strength (*r* = 0.24, p<0.001), despite all resulting in near-complete splicing. While few weak splice sites were included in the analysis, the observed trend suggests that introns with weak 5SS motifs may exert little or no IME. No analogous relationship was observed between 3SS strength and IME (*r* = –0.04, p=0.42; Fig. 3c, Supplementary Fig. 3d).

The dependence of IME on 5SS strength suggests that IME may depend on factors specific to 5SS recognition. For major introns, this involves U1 snRNP and subsequently U6 snRNP and PRP8, while for minor introns (Fig. 3a) the U11 and U6atac snRNPs play roles analogous to U1 and U6, respectively^16^. Replacing the native major splice sites with strong P120 minor splice sites yielded mostly negative or zero IFC values even when the intron remained well-spliced, suggesting that the minor spliceosome may be incapable of eliciting IME (Fig. 3d,e). Previously, it has been reported that the splicing of minor introns can be rate-limiting for gene expression^20^ and that transcripts containing unspliced minor introns may be subject to nuclear RNA decay^21^, potentially contributing to this phenomenon.

### A second screen assesses impact of minor intron bodies and diverse SS motifs

Our screen of natural introns uncovered not only introns that substantially enhance gene expression (Fig. 3f) and elucidated contributions of the intron body, but also suggested that the 5SS motif may be a key determinant of IME. However, these observations were based on just a few dozen moderate to strong 5SS motifs and a handful of weak 5SS motifs, since stronger 5SS are more common in natural introns and the exonic portions of splice sites were held fixed in our library to match the intronless controls. Further, we only considered a single, strong minor splice site pair and only one minor intron body, limiting what we could conclude about the effects of the minor spliceosome.

Therefore, to validate key results of this screen and address these issues more systematically, we designed a second library encompassing all 3,386 5SS motifs used in human genes (positions –3 to +6) (Fig. 4a). To assess the effect of each human 5SS motif on IME – while controlling for intron body contributions – we selected one human intron containing that motif and replaced the intron body with three fixed major intron body sequences (*WASH7P* i10, *CD1C* i5, *MKNK1-AS1* i7) and one minor intron body (*STX10* i2). The three major introns, all present in the first screen, were chosen because they were well-spliced, lacked restriction sites, and varied greatly in sequence composition, though the *MKNK1-AS1* intron required internal truncation to fit the tighter length constraints of this library design. *STX10* i2 as a representative minor intron because initial qPCR experiment suggested it could exhibit IME with major splice sites (Supplementary Fig. 4a). To further assess whether minor intron bodies are compatible with IME, we also included a diverse collection of minor introns, flanked by their native SS or by strong major (UbC) splice sites. To eliminate potential amplification bias arising from differences in splice junction sequences, we included separate intronless controls for each unique exon-exon junction sequence in the library (Fig. 4a). As with the first screen, each intron was associated with three distinct barcodes, with IFC averaged between the three values. Sequences, PS and IFC values, etc., for individual introns in the second screen are listed in Supplementary Table S5.

**Figure 4:**
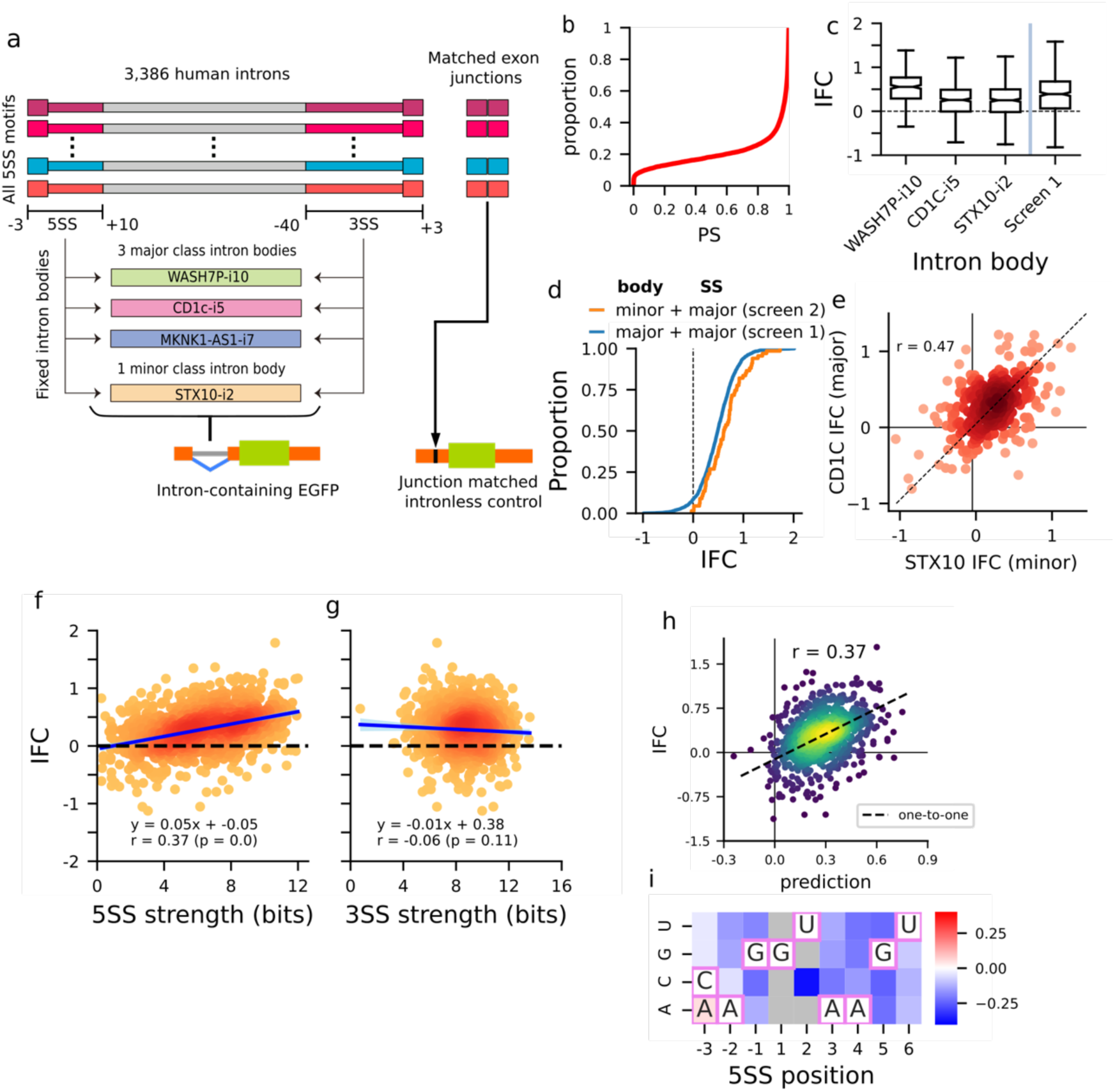
Assaying the enhancing effects of all human 5SS motifs. **a)** The design of the second intron library. Splice sites from 3,386 human introns, spanning all human 5SS motifs, were coupled with four fixed intron bodies and inserted into EGFP reporters, as with the first screen, along with junction matched intronless controls. **b)** Cumulative distribution of proportion spliced. **c)** The distributions of IFC for the different intron bodies, compared to those observed in the natural intron library. **d)** The distribution of IFC for minor intron bodies and major intron bodies coupled to strong major splice sites. **e)** Correlation of the effects of major splice sites on IFC when coupled to minor (*STX10*) or major (*CD1C*) intron bodies. **f, g)** The association between mean IFC (averaged across the well-spliced intron bodies) and strength of the 5SS (**f**) or 3SS (**g**). Trend lines are linear regression fits, with 94%-intervals constructed by bootstrapping splice sites 10,000 times. **h)** The predicted versus observed effect of 5SS sequences on expression based on the linear model of 5SS motif composition. **i)** Heat map shows inferred effects of non-consensus nucleotides on the magnitude of IME (all negative); gray indicates bases observed in fewer than ten 5SS motifs. Consensus bases are shown, boxed in pink.

The resulting library yielded a wider range of splicing variation than observed in the first screen, as expected from the broader range of splice site strengths (Fig. 4b). The assayed splice site combinations yielded consistent PS values across three of the four intron bodies (Supplementary Fig. 4b). The exception was the modified *MKNK1-AS1* i7 intron body which exhibited high rates of cryptic splice site usage (Supplementary Fig. 4c); we therefore excluded it from our downstream analyses and focused on splice site pairs with <1% cryptic splicing. PS was positively correlated with the strength of both the 5SS and 3SS, as expected (Supplementary Fig. 4d). As in the first screen, intron-containing EGFPs were generally more highly expressed than their matched intronless controls (Supplementary Fig. 4e), with the range of IFC for well-spliced introns comparable to that observed in the first screen (Fig. 4c).

Minor introns in this library were often poorly spliced when coupled to their native splice sites – especially those with AT-AC splice site dinucleotides, suggesting frequent dependence of their splicing on additional contextual features (e.g., exon sequences, splicing of flanking introns). However, replacing their splice sites with the strong UbC major splice sites yielded IME comparable to that observed with major intron bodies, both in our library (Fig. 4d) and in additional minigene experiments (Supplementary Fig. 4a). Further, major splice site pairs elicited IME when coupled to the minor *STX10* i2 body, comparable to the levels observed when paired to major intron bodies (Fig. 4e), suggesting that minor intron bodies are compatible with IME driven by the major spliceosome. Minor introns have previously been reported to reduce gene expression, likely reflecting nuclear decay of slowly spliced transcripts^21^. Consistent with this possibility, comparably reduced expression was also observed when minor splice sites were replaced with those of beta-globin intron 1 (Supplementary Fig. 4f). Beta-globin i1 has been previously reported to undergo slow splicing^14^ and was one of the few major introns found to decrease expression in the first screen.

### Confirmation that strength of the 5’SS, but not the 3’SS, is an important determinant of IME

In all, 821 splice site pairs were well-spliced in at least one of the intron bodies. This included 184 weak, 389 moderate, and 248 strong 5SS (up from fewer than 50 of each in the natural intron screen). The range of IFC for well-spliced introns was consistent with that observed in the first screen (Fig. 4c). Consistent with our finding above that intron body impacts IME, median IFC differed among the intron bodies considered in this screen (*WASH7P* i10: 0.55, *CD1C* i5: 0.26, *STX10* i2 0.25), with *MKNK1-AS1* having too few well-spliced introns to evaluate IME. As suggested by the natural intron screen, 5SS strength was associated with IME across these 821 5SS motifs (*r* = 0.37, Fig. 4f). Strong 5SS elicited the most potent IME, with weaker motifs being associated with weaker IME, and the weakest 5SS eliciting little to no IME, despite all being well-spliced. No association with 3SS strength was evident (*r* = –0.06, Fig. 4g).

The number of distinct 5SS motifs in this screen provided an opportunity to more directly investigate which 5SS features contribute to IME. To ask how different consensus and non-consensus bases in a 5SS motif impact IME, we used a linear model to predict IFC based on the sequence composition of the associated 5SS motif. This approach predicts IFC as well as MaxEnt-based splice site strength (Fig. 4h), and predicts that the canonical 5SS consensus MAG/GUAAGU (M=A or C) – which is recognized by the complementary motif in U1 snRNA – is the most IME-inducing motif (expected IFC 0.7). Every non-consensus nucleotide at positions –2 to +6 of a 5SS is predicted to reduce IME (Fig. 4i), with the strongest reductions inferred for positions –1 to +5. These data confirm that every consensus position of the 5SS contributes to IME, and indicate that 5SS strength and IME are intimately related.

### Natural variation in first 5SS impacts gene expression in a pattern consistent with IME

The observed association between 5SS strength and IME suggests that mutations in splice sites might alter the magnitude of IME even when they do not impact splicing. To investigate this possibility, we used the set of quantitative trait loci (QTL) identified by the GTEx consortium^22^. These data provide both genetic and expression variation across hundreds of mostly healthy individuals, spanning dozens of tissues, allowing the identification of genetic variants associated with expression variation (eQTLs) and splicing variation (sQTLs). We focused on eQTLs that were located within a 5SS or 3SS motif and asked whether variants that alter splice site strength without affecting splicing were associated with altered expression in a pattern consistent with changes in IME. To control for potential effects of altered splicing on transcript stability, we excluded any eQTL that was also associated with splicing variation. IME is known to be dependent on intron position within a gene, with the strongest effects observed for introns located most proximally to the TSS, motivating us to focus on first introns versus later introns in transcripts. So that intron numbers could be unambiguously assigned, we focused on eQTLs in canonical MANE-select transcripts and asked whether the observed associations depended on the intron’s location in the transcript. We found that 5SS variants–but not 3SS variants–associated with gene expression variation were enriched in the first intron of genes (Fig. 5a, b). The effects of these first 5SS eQTLs on expression were substantially stronger than eQTLs in the 5SS of other introns or in 3SS in any intron (Fig. 5c). Further, the effect of a first-5SS eQTL on expression related to the predicted effect of that variant on the splice site’s strength: variants that strengthened or weakened the splice site were associated with increased and decreased gene expression, respectively (Fig. 5d). This association was specific to eQTLs in the first 5SS, with variants in non-first 5SS (Fig. 5e) or in 3SS (Fig. 5f,g) displaying no relationship between changes in splice site strength and associated expression change. Together, these observations suggest that the effects of 5SS motifs on gene expression observed in our screens shape human gene expression variation in natural populations and that variation in promoter-proximal 5SS impacts gene expression by modulating IME.

**Figure 5:**
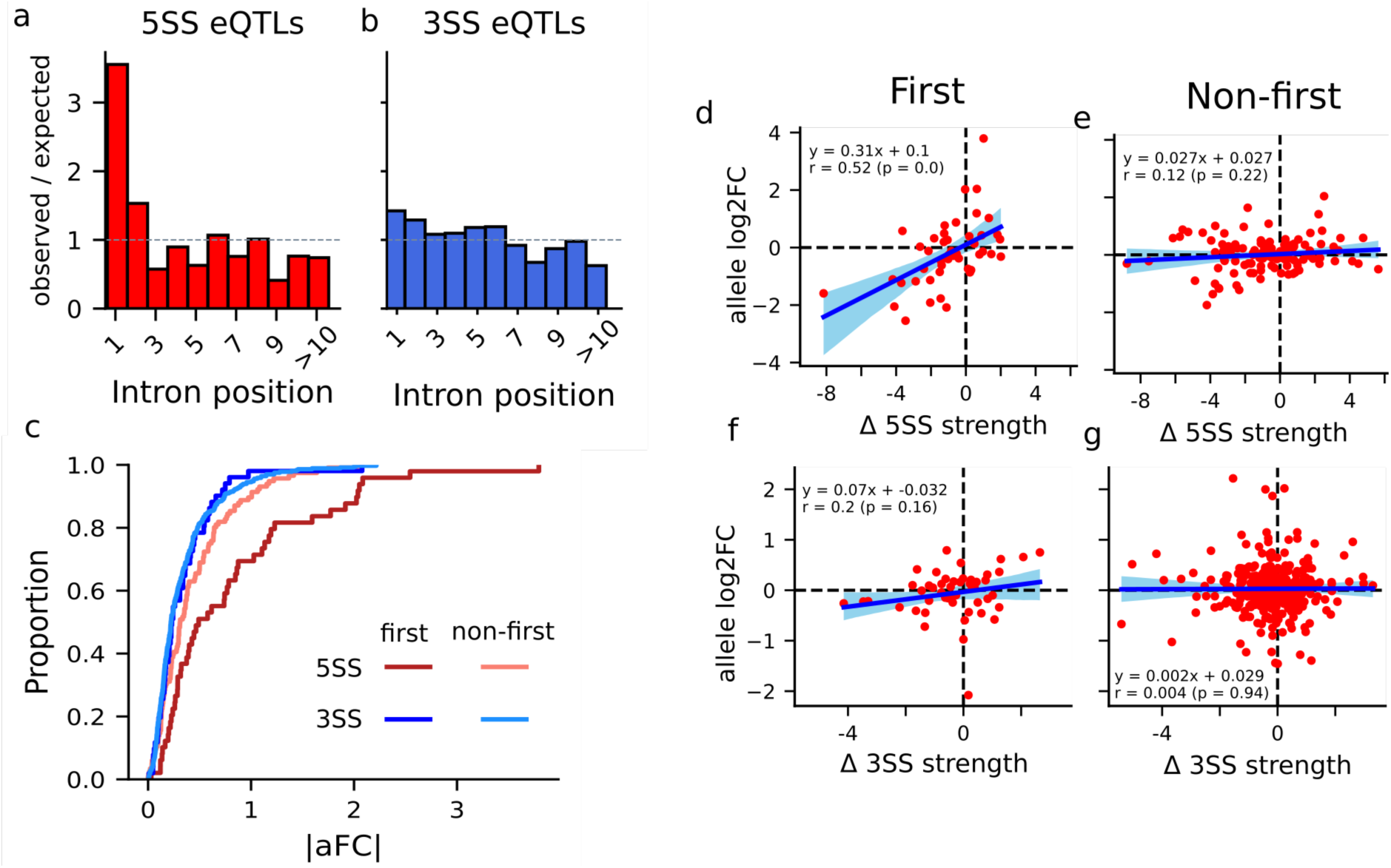
Association between natural variation at human splice sites and gene expression. **a, b**) Enrichment of 5SS (a) and 3SS (b) across intron order relative to a uniform expectation. **c**) ECDFs of the absolute magnitude of the effect sizes on gene expression of 5SS (red) and 3SS (blue) eQTLs in first (dark) and non-first (light) introns. **d-g**) The relationship between the predicted effect of an eQTL on splice site strength and its effect on gene expression for first 5SSs (d), non-first 5SSs (e), first 3SSs (f), and non-first 3SSs (g); Trend lines are linear regression fits, with 94%-intervals constructed by bootstrap sampling eQTLs 10,000 times.

### Perturbations to splicing machinery impact gene expression through IME

Our findings regarding the impact of 5SS strength on IME suggest that perturbations to U1 snRNP that alter 5SS recognition, such as those found in certain hematologic malignancies ^23 24^, might also have direct effects on gene expression via changes in the potency of IME. One particularly striking example is a recurrent mutation in U1 snRNA (U1^g.3A>C^) at the position which pairs with the +6 position of 5SS motifs. This change in specificity from the canonical +6U to +6G (Fig. 6a) was reported to result in broad splicing changes in chronic lymphocytic leukemia (CLL) patients ^23^. In addition to altered splicing, expression changes were also reported in hundreds of genes. While some of this altered expression may reflect alterations in isoforms with distinct stabilities or secondary effects, we reasoned that the altered 5SS specificity might also directly impact gene expression by modulating IME at specific loci.

**Figure 6:**
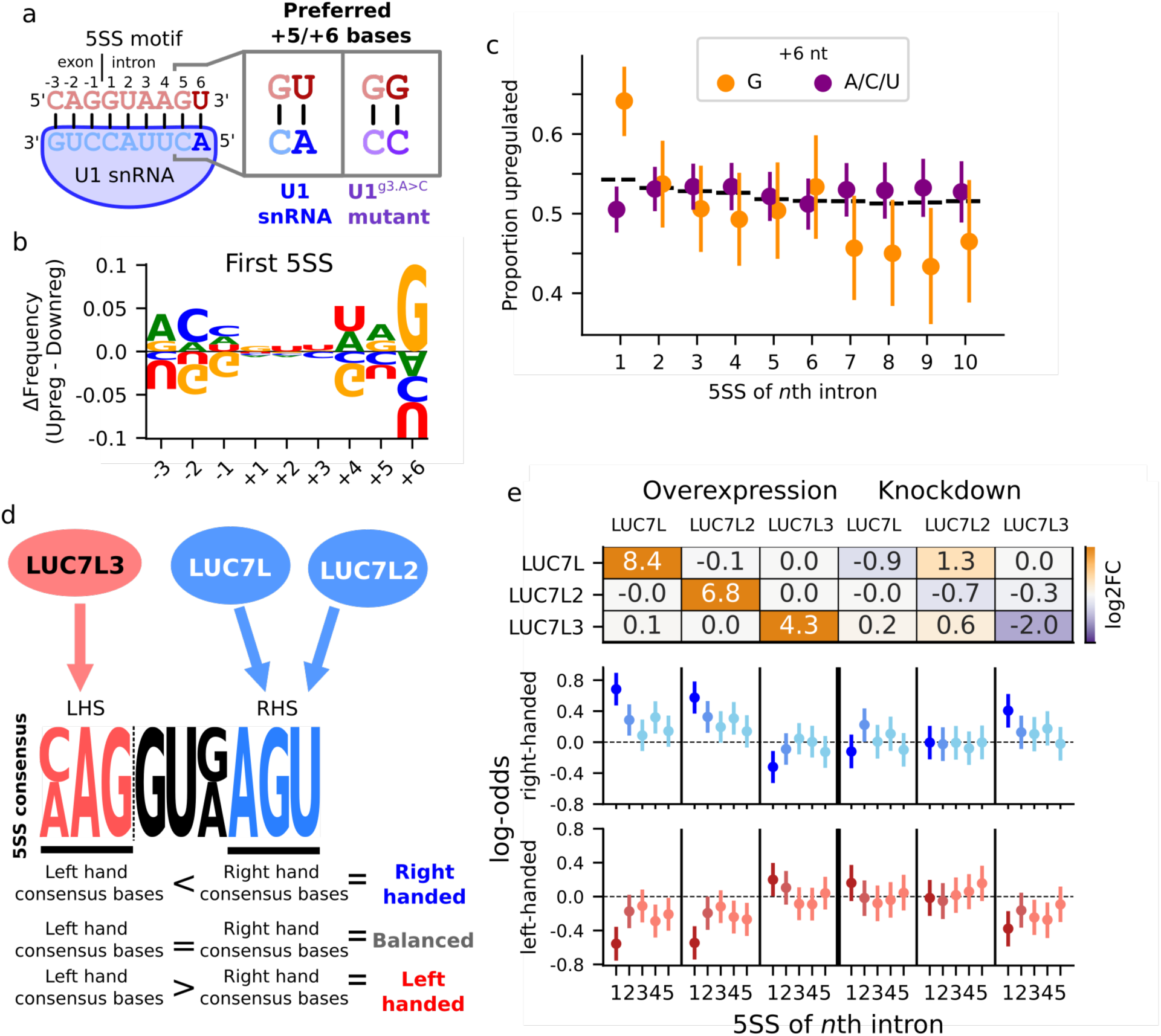
Effects of U1 snRNP perturbations on gene expression. a) Schematic showing complementarity between the 5’ end of U1 snRNA and the canonical 5SS motif and how specificity for the +6 position is altered by the U1^3g.A>C^ mutation. b) The difference in nucleotide frequencies in the first 5SS motif of genes upregulated in CLL patients with the U1^3g.A>C^ mutation versus genes that are downregulated. c) The frequency of upregulation among significantly differential expressed genes with or without a +6G at a given 5SS motif, with 94% credible intervals computed under a beta binomial conjugate model. The +6 position of only the first 5SS is associated with upregulation. d) Overview of how proteins in the LUC7 family modulate 5SS motif preference, with LUC7L and LUC7L2 favoring 5SSs with more consensus bases on the intronic side (right-handed) and LUC7L3 favoring 5SSs with more consensus bases on the exonic side (left-handed). e) Top: the changes in expression of LUC7 family genes upon overexpression or knockdown of a particular paralog. Bottom: The log2-odds enrichment of right-handed and left-handed motifs across the first five splice sites of genes in the top and bottom quintiles of differential expression in LUC7 protein in these perturbation data, with 94% credible intervals computed under a beta binomial conjugate model with uniform priors.

We reasoned that, if IME contributes to these expression changes, the presence of a +6G in a gene’s first 5SS should be associated with up-regulation. To assess this possibility, we examined the genes reported as differentially expressed (10% FDR) in samples from CLL patients with the U1^g.3A>C^ mutation^23^. Consistent with altered IME, +6G was enriched in the first 5SS motifs of up-regulated genes compared to those of down-regulated genes (Fig. 6b, Supplementary Fig. 5a,b). As a control, we examined non-first 5SS of up- and down-regulated genes, and observed no such enrichment for +6G. Among differentially expressed genes, those with +6G in their first 5SS were more likely to be upregulated (64%) than those with +6A/C/U (50%), while the presence of a +6G in non-first splice sites was not associated with up-regulation (Fig. 6c). The specificity of this effect for the first 5SS is consistent with IME, rather than potential effects of aberrant splicing on message stability, which would be associated with splice site motifs across all intron positions. Based on the thermodynamics of RNA folding, it is anticipated that basepairing at the +6 position would likely be insufficient to compensate for a +5 bulge. Consistent with this hypothesis, the effects of +6G on IME appear specific to 5SS motifs with the canonical +5G (Supplementary Fig. 5c). Perturbations of other U1 or U6 snRNP proteins and U1 snRNP auxiliary factors may also impact

5SS recognition. We recently reported that the U1-associated Luc7-like proteins impact 5SS choice in a manner dependent on the 5SS motif^25^. Paralogs LUC7L and LUC7L2 favor motifs with consensus bases on the intronic side (“right-handed” motifs, RH), while LUC7L3 favors the usage of 5SS motifs with consensus bases on the exonic side (“left-handed”, LH motifs) (Fig. 6d). Re-examining overexpression and knockdown data for these factors, we observed that gene expression changes upon LUC7 protein perturbation are associated with the pattern of consensus matching (“handedness”) of the first 5SS (Fig. 6e). RH motifs are enriched at the first 5SS of genes up-regulated upon *LUC7L*/*LUC7L2* overexpression and depleted in genes up-regulated upon *LUC7L3* overexpression. The opposite effects are observed upon *LUC7L* and *LUC7L3* knockdown. No significant effects were observed in the *LUC7L2* knockdown – potentially due to compensation by *LUC7L* (Fig. 6e, top) via a previously described regulatory mechanism^25^. The associations we report between perturbations to U1 and its auxiliary factors and expression variation, dependent on a gene’s first 5SS motif, suggest that altering U1’s 5SS specificity leads to broad gene expression changes via modulation of IME.

## Discussion

Here we screened thousands of natural introns and 3SS motifs, and all human 5SS motifs to determine rules for how intron sequence impacts gene expression. Surprisingly, we find that whether an intron induces IME depends upon the strength of its 5SS – but not its 3SS – requiring at least a moderate strength motif (∼4 bits or stronger), with the largest effects observed for the strongest 5SS motifs. Since we restricted our analyses to introns with near-complete splicing, this variation does not reflect differences in splicing outcome but instead suggests that the kinetics of splicing or of snRNP binding to the 5SS is a key driver of IME. We show that the sequences of intron bodies also impact expression, with U-rich motifs having the strongest positive effects, consistent with our previous work^15^. IME in this system appears specific to the major spliceosome, as replacing major splice sites with minor splice sites consistently eliminated IME, even when splicing remained efficient. Moreover, replacing an intron’s natural minor splice sites with strong major splice sites was sufficient to confer IME. Short introns are often used in gene therapy or synthetic biology applications to boost expression^26,27^. Our findings emphasize the importance of 5SS strength, intron body features and splicing by the major spliceosome in optimizing design of introns with potential to modulate expression.

While our study focused on the regulatory rules governing IME, some of our findings implicate specific splicing machinery. The fact that consensus matching at all positions on both the exonic and intronic sides of the 5SS motif contribute to IME (Fig. 4i) is more consistent with involvement of U1 snRNP – which interacts across the whole 9-nt motif – than U6 snRNP, which interacts with only a few positions on the intronic side of the motif^18^. Recruitment of U1 snRNP might positively impact the expression of a gene by enhancing transcription initiation or elongation or by inhibiting premature termination. An early study reported that mutations disrupting 5SS in two genes reduced their expression, potentially by affecting recruitment of transcription initiation factors to the promoter^4^. Work in budding yeast suggests that this may be facilitated by gene looping, such that the promoter and 5SS are in close physical proximity^9,28^. Another possibility is that assembly of splicing machinery including U1 and U2 may promote productive PolII elongation^5^. Further, recruitment of U1 snRNP may inhibit transcription termination via the canonical cleavage and polyadenylation (CPA) pathway as U1 snRNP can inhibit use of nearby downstream polyadenylation signals (PAS)^29^. However, none of the intron bodies considered in our splice site library contained potential PAS motifs (Methods), suggesting that other mechanisms are active in our system. Alternatively, a recent structure of U1 snRNP in complex with RNAPII and nascent RNA shows that U1 associates with the same interface of RNAPII as the integrator pre-termination complex – which drives promoter-proximal termination independent of CPA – raising the possibility that binding of U1 snRNP near the TSS may shield the transcript from integrator-mediated termination^30^. Of course, effects on other steps in gene expression may also contribute to IME observed in our reporter and endogenously^1^.

Our results indicate that a gene’s first 5SS plays a dual role, impacting gene expression as well as splicing. This finding has implications for interpretation of mutations in splice sites and the splicing machinery; in both cases phenotypic effects are typically assumed to be mediated by their effects on splicing^31^. In the context of human genetics, we found evidence that variants can alter IME by strengthening or weakening a gene’s first 5SS without necessarily altering splicing, representing an important new category of eQTLs. We also show that U1 snRNA mutations in CLL upregulate the expression of genes whose first 5SS matches the newly favored motif (Fig. 6c) indicate that alterations to trans-acting factors may also exert their effects by directly impacting gene expression rather than splicing. Similar effects are likely to occur in other U1 snRNA-associated cancers, e.g., 97% of adult sonic hedgehog medulloblastomas have U1 snRNA mutations^32^, and potentially also for mutations to protein-coding splicing factors. For example, we find evidence that the U1 auxiliary factor *LUC7L2* – commonly deleted in AML – exerts similar, 5SS motif-dependent effects on expression (Fig. 6e). Such 5SS-based expression regulation is likely to extend to a range of other spliceosomopathies^31^. Beyond mutation interpretation, our findings further suggest that modulating the recognition of a gene’s first 5SS may be an important aspect of physiological gene expression programs, and is a potential avenue for boosting the expression of specific genes in synthetic or therapeutic contexts.

## METHODS

### Design of first screen

The list of annotated introns was downloaded from the UCSC table browser^33^. Intron sequences were retrieved from NCBI RefSeq. Gene expression data for K562, HEK293 and HepG2 cells reported by Ramskold and coworkers were used^34^. Introns shorter than 190 nt were classified by feature types, and a set of randomly selected introns from each group was used for library construction. In addition to their native splice sites, we also replaced the first 12 and last 40 nt of each intron with strong major splice sites (the corresponding regions of *UbC* i1 and *EF1A1* i1, respectively) and strong minor splice sites (sequences from *P120* i7). The EF1A1 3SS was used rather than the UbC 3SS because it has a longer downstream 5’ UTR sequence naturally, providing a more extended natural exonic context and space between intron, barcode and EGFP coding sequence. Non-intronic regions from 5’ UTR, 3’ UTR, and coding sequences were randomly selected as 190-nt fragments and attached to both U2- and U12-type splice sites. We included ten intronless controls with distinct barcodes. We additionally added control constructs with only a 5SS or only a 3SS, which both resulted in reduced EGFP detection. However, these observations are difficult to interpret as they might result from cryptic splicing involving sequences beyond the ampliconic region or reduced stability of transcripts with partially assembled splicing machinery and were not considered further. Sequences containing ATG codons or restriction enzyme sites used for cloning (BbsI, BsaI, Esp3I) within the intron were mutated, with compensatory mutations used avoid altering the intron’s overall base composition. Each intron sequence was paired with a unique 12-nt barcode and flanking primer binding sites, as previously described^15^. Oligos shorter than 300 nt were synthesized as DNA oligo pools, while longer fragments were synthesized as individual DNA fragments (Twist Bioscience, CA, USA). We used identical UbC promoters, 3’ UTRs and polyadenylation signals for the EGFP and dTom reporters, eliminating small differences in the 3’ UTRs present in our previous screen.

### Design of second screen

The –3 to +10 of the 5’SS and –40 to +3 of the 3’SS were extracted from the intron sequences obtained from the USCS table browser and RefSeq as described above. Sequences containing restriction enzyme sites used for cloning in the splice sites were excluded. All unique –3 to +6 regions of the 5’ splice site were included; if multiple introns shared the same sequence, the intron corresponding to the most highly expressed mRNA in HEK293 cells was chosen. Selected splice sites were ligated to each of four intron bodies (W*ASH7P* i10, *CD1C* i5, *MKNK1-AS1* i7, *STX10* i2). The intron bodies were examined to confirm the absence of any polyadenylation signals hexamers (AAUAAA or AUUAAA), which were not present in any of the introns.

The list of minor (U12-type) introns was obtained from the U12 Intron Database (U12DB)^35^. The U12-type introns shorter than 151 nt that lacked the restriction enzyme sites used for cloning (BbsI, BsaI and Esp3I) were selected as introns to test. One 100-nt fragment was randomly selected from the 128 randomly-chosen U12-type introns longer than 152-nt and its cognate, UbC or beta-globin splice site were attached to create the fragment U12-type intron constructs. All unique junction sequences were cataloged, and junction-only (intronless) controls were included in the library design.

The 12nt barcodes were randomly generated in a manner to avoid containing an ATG, a restriction enzyme site (BbsI, BsaI and Esp3I) used in cloning, a run of G longer than 3-nt, an A, C or T run longer than 4-nt, more than 6 Gs, or more than 8 nt of A, C or T. Each designed intron or control was randomly paired to a barcode, such that each new barcode had a Levenshtein edit distance ≥ 3 (representing at least 3 insertions/deletions/substitutions) from any previously used barcode to minimize barcode mis-mapping.

### Cloning

The primers and plasmids used for cloning are listed on Supplementary Tables S1-S3. The natural intron library was constructed by PCR amplification of the oligo DNA pool, followed by cloning of the PCR products into plasmid pMBYS124, as previously described^15^. The natural splice site library was constructed using a similar approach, except libraries were prepared separately by oligo length (>250 nt or ≤250 nt) to minimize cloning bias. Plasmid backbone pMBYS257 and dTom cassette plasmid pMBYS258 was used for cloning instead of pMBYS124 and pMBYS126, respectively. These plasmids have the splice junction sequence in the natural intron library removed, allowing for matched splice junctions for both dTom and EGFP mRNA when transcribed and spliced.

### Long-read sequencing and analysis

Plasmid libraries were linearized with one of the following restriction enzymes: XmnI, NruI, SspI, or ScaI, and subsequently purified using 0.6× SPRI beads (Beckman). Sequencing was performed on an Oxford Nanopore PromethION at the MIT BioMicroCenter. The Minimap2 algorithm^36^ was used to align the reads to the sequence mimicking the plasmid library with barcodes and intron regions marked as Ns. BAM files were processed with Samtools^37^. The sequence aligned to the barcodes of the reference were extracted and matched the list of the barcode sequences present in the sequence library. Since Nanopore’s long read sequencing has moderately high mutation rates, and because the barcodes were designed to differ from each other by at least three nucleotides, up to one mismatch was allowed to count the barcodes detected in the reads. Barcode counts for each construct were calculated.

### Cell culture and transfection

HEK293T A2 and HeLa A12 HILO-RMCE cells were kindly provided by Eugene V. Makeyev^38^. The cells were grown and transfected as described previously^15^. For the natural splice site library, two sub-libraries—one comprising oligos shorter than 250 nt and the other comprising oligos longer than 250 nt—were mixed at a 1:8 ratio reflecting the relative number of constructs in each sub-library prior to transfection. For the RT-qPCR experiment, 1.5×10^5^ cells were transfected with using Lipofectamine 3000 (Thermo).

### Reverse transcription and qPCR for assessing IME with specific minor intron bodies

Total RNA was extracted from cells using the Quick-RNA MiniPrep Kit (Zymo Research) with on-column DNase I treatment. The purified RNA was quantified using a NanoDrop spectrophotometer, and 300 ng of RNA was reverse transcribed using SuperScript IV VILO Master Mix (Thermo). The resulting cDNA was used as a template for qPCR. The mRNA levels of EGFP, dTomato, and GAPDH were quantified by qPCR using gene-specific primers and probes. qPCR reactions were performed on a QuantStudio 5 Real-Time PCR System (Thermo) according to the manufacturer’s instructions. For each target (EGFP, dTomato, and GAPDH), a standard curve was generated using serial dilutions of a plasmid with known copy number, and the copy number of each transcript in the samples was calculated based on the corresponding standard curve. The mRNA levels of EGFP and dTomato were normalized to those of GAPDH, which served as an internal control, and the relative expression levels were calculated accordingly. All reactions were performed in technical triplicate, and the mean values were used for subsequent analysis.

### RNA-seq preparation and sequencing run

After 48-hours of transfection, the cells were trypsinized and the total RNA was extracted using TRI-reagent and Direct-zol RNA Miniprep Plus kit (Zymo Research). The residual DNA was removed by treating the total RNA with ezDNase (Thermo) followed by heat treatment to inactivate the activity. EGFP and dTom mRNAs were reverse-transcribed to cDNA using SuperScript IV (Thermo) with a mixture of gene-specific primers. cDNA was treated with RNase cocktail, and sequencing libraries were generated by PCR (12 cycles for HEK293, 15 cycles for HeLa). Primers targeting a common region of both genes and incorporating sample indexes were used. The amplicon was purified using 2.5x volume of SPRI beads (Beckman). The sequencing was performed with 150-bp paired-end reads using an Element AVITI instrument at the MIT BioMicroCenter.

### Data analysis of natural intron library and the natural splice site library

Custom Python scripts were used for sequencing data analysis. Only reads containing the correct index sequence were used for analysis. For the natural intron library, the first 8 nucleotides of each read generated using sequencing primer YS331F were examined to classify the transcripts as EGFP or dTom mRNA. Next, reads covering the upstream region of the barcode were scanned to identify spliced EGFP reads based on the presence of the expected splice junction sequence. Among the remaining reads, unique 8- or 7-nucleotide sequences (designed frames for EGFP transcripts) were detected to further classify EGFP reads as spliced or unspliced. The corresponding 12-nt barcode sequences were extracted from classified reads and counted. Barcode counts were normalized using DESeq2, and log_2_ fold-change values were calculated. Only barcodes detected in all transfection replicates with at least 100 reads supporting each dTom and EGFP were retained for downstream analyses.

### Estimating intron-associated fold-change

We used DeSeq2 to evaluate the log-fold difference in expression between EGFP and dTom for each barcode across replicates^39^. Briefly, we computed size factors that account for baseline differences in dTom versus EGFP expression, using the intronless constructs as controls. We then estimated the log_2_ fold change (LFC) associated with each intron-barcode pair using a paired design `∼replicate + isGFP` where `replicate` represents the replicate-specific dTom expression level and `isGFP` reflects the difference in GFP and dTom expression across all replicates. Most intron sequences in our library are coupled to 3 barcodes so that we can disentangle the effect of the intron from the potential effect of the barcode on GFP expression. We estimate this by averaging across the *k* (typically 3) barcodes:

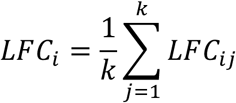

and estimate the associated uncertainty of this average based on the uncertainty associated with each of the individual log_2_ fold-changes

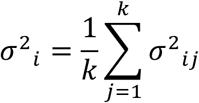

To estimate the intron-associated log_2_ fold-change (IFC), we normalize to the average log-fold change associated with the corresponding intronless control

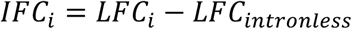

and compute the uncertainty as the sum of the two variances

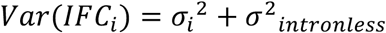

In our first screen, the intronless control values are simply the average across all intronless constructs. In our second screen, it is the average across all exon junction-matched intronless controls. To evaluate significantly enhanced or repressed EGFP expression relative to these intronless controls we employed ashr^40^, using a half-normal mixture, to accomplish a multiple-testing correction of the effect size estimates. We defined significance at a 10% false sign rate.

### Estimating proportion spliced

We compute the proportion of spliced reads (PS) for each intron-barcode pair from the number of observed spliced reads, S, and unspliced reads, U:

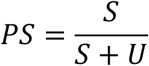

When choosing a PS cutoff for identifying well-spliced introns, we considered the potential for underrepresentation of the longer unspliced products in amplicon amplification and sequencing^41^. Calibrating PS using an estimate of this potential underrepresentation derived from predominantly unspliced (low PS) introns indicated that a cutoff of PS≥0.99 represents at least 90% intron excision at a minimum.

### Scoring splicing signals and regulatory elements

We quantify splice site strength using the updated, third-order MaxEntScan models from SMsplice^17^, and similarly employ the SMsplice SRE hexamer scores to quantify intronic cis-elements. To compare these hexamer scores to the tetramer enrichments observed in our screens, we assign each tetramer the average SRE score of all hexamers in which it occurs.

### Regression analyses and uncertainty quantification

We quantified the relationships between splice site strength and gene expression using ordinary least squares linear regression (scipy.stats.linregress). To estimate uncertainty while accounting for the potential influence of outliers, we bootstrapped the data in 10,000 replicates. To assess the impacts of non-consensus bases in 5SS motifs on IME, we employed a linear model (statsmodels.OLS) to predict the measured IFC of each well-spliced construct based on the sequence composition of its 5SS. The intercept of the model represents the expected IFC of a consensus motif and the regression coefficients represent the effects of each possible non-consensus base at positions –3 to +6. To assess the sub-additive relationship observed between U_4_ motifs and IFC, we fit a generalized sigmoid with the steepness, midpoint, and asymptotes fit to the observations by minimizing the squared error (scipy.optimize.minimize).

### eQTL analyses

GTEx cis-eQTLs and cis-sQTLs were downloaded from the GTEx portal (v.11 release) at https://www.gtexportal.org/home/downloads/adult-gtex/qtl (filenames:GTEx_Analysis_v11_eQTL.tar and GTEx_Analysis_v11_sQTL.tar). For each tissue, the *.egenes.txt.gz file was filtered for the most significant eVariant. Using bedtools window -sm-sw, 5SS eQTLs were defined as eVariants within 3 nt upstream and 5 nt downstream of intron start positions. Similarly, 3SS eQTLs were defined as eVariants within 16 nt upstream and 1 nt downstream of intron end positions. These windows were chosen so that the search space for 3SS eQTLs was double of that for 5SS eQTLs. Intron start and end positions were defined based on the GENCODE v47 primary assembly basic annotation. Splice site strength of 5SS and 3SS eQTLs were scored with an updated version^17^ of MaxEntScan that captures dependencies between triplets of positions. 5SS and 3SS eQTLs that overlapped significant sQTLs with *q*≤0.05 reported in *.sgenes.txt.gz were excluded from our analyses. MANE Select isoforms from the GENCODE v47 primary assembly basic annotation were used to unambiguously assign an intron number (based on position in the gene) to 5SS and 3SS eQTLs. 5SS and 3SS eQTLs located near splice sites that are absent from MANE Select isoforms were excluded from analyses based on intron number.

### Alignment and analysis of LUC7 knockdown/overexpression data

We downloaded *LUC7L*, *LUC7L2*, and *LUC7L3* knockdown (E-MTAB-9709) and overexpression (PRJNA1200827) RNAseq dataset from SRA and aligned to the hg38 human reference genome with STAR2, using the GENCODE v38 genome annotations and two-pass alignment. We quantified differential gene expression using DESEQ2 to estimate log2FC and associated posterior standard deviations for each gene. To account for potential sequence biases, we prepared the DESEQ inputs using Salmon with the --gcBias flag and corrected any remaining biases by regressing each gene’s estimated log2FCs against %GC and its square, similar to the approach used in ENCODE^42^. To investigate general patterns of differential expression without significance filtering, we quantified the signal-to-noise ratio of each estimated effect size as a z-score, normalizing the gene’s logFC to the associated posterior standard deviation. We filtered out genes averaging fewer than 100 normalized read counts across samples and selected genes in the top and bottom quintiles of z-scores to assess whether genes whose expression increased or decreased upon perturbation had characteristic 5SS motifs.

## Data availability

The read counts used to estimate IFC, filters used in our analyses, and information about the constructs associated with each barcode (e.g. splice site strength, intron sequence) provided in Supplementary Tables S4 and S5, for screens 1 and 2 respectively.

## Acknowledgements

We thank Mikko Frilander, Phillip Sharp and members of the Burge lab for helpful comments on this manuscript. This work was support by NIH grant 5-R01-HG002439 (C.B.B.) and by a HEALS grant from MIT (C.B.B.).

---

**Supplementary figure 1:**
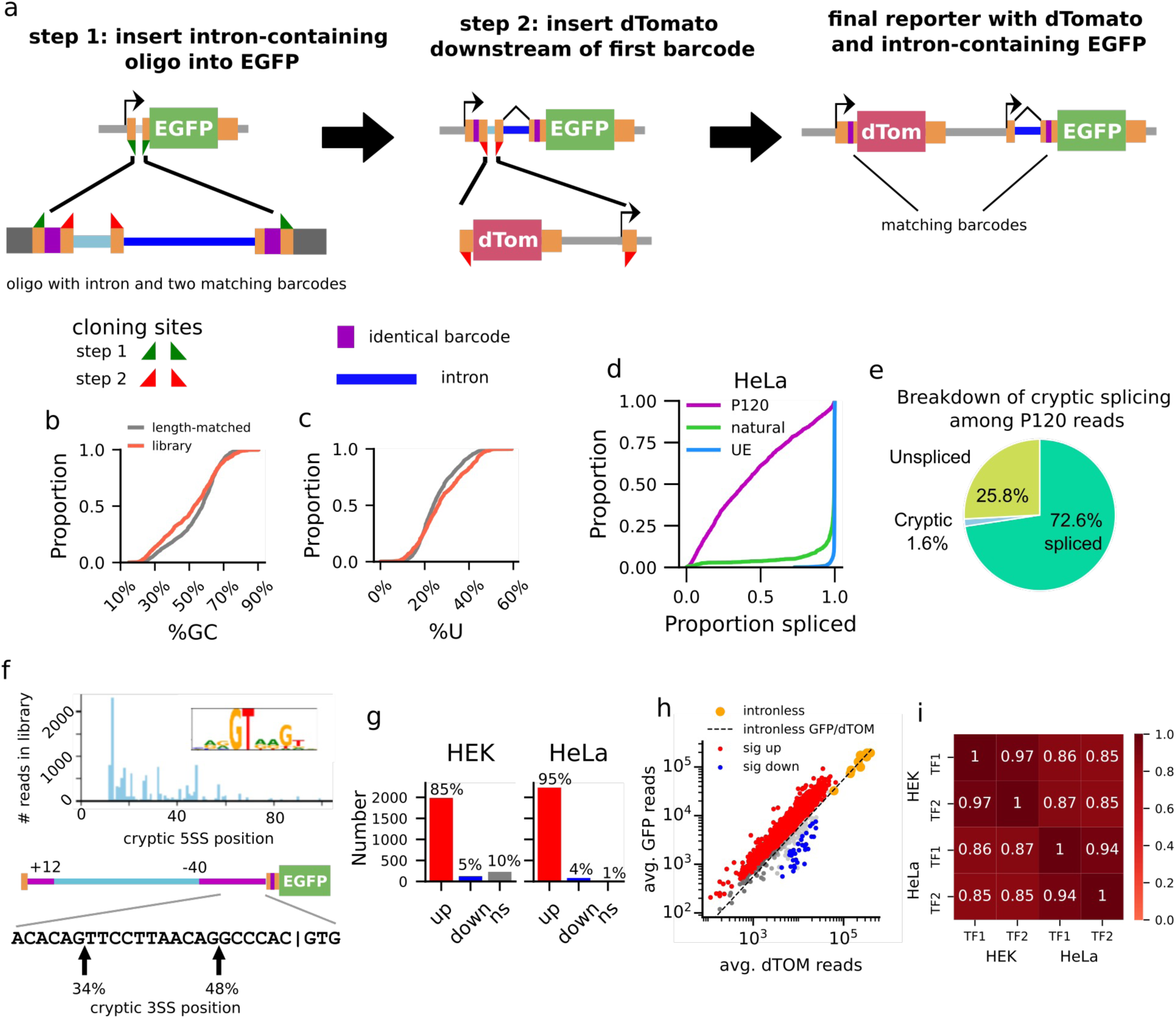
Cloning strategy and features of initial screen. **a)** Overview of the two-step cloning strategy used to ensure that the intron-containing EGFP and intronless dTomato in each construct have matching, intron-specific barcodes. **b, c)** The distribution of %GC (**b**) and %U (**c**) for the screened natural introns, with length-matched introns selected from the genome annotation included for comparison. **d)** The proportion spliced of natural introns in HeLa with their native splice sites, strong major splice sites (UE), and strong minor splice sites (P120). **e)** The proportion of P120 EGFP reads in the library spliced as expected, unspliced, or cryptically spliced **f)** Overview of cryptic splice site positions. Top: the distribution and consensus motif of cryptic 5SSs. Bottom: The positions of cryptic major 3SSs in the P120 3SS. **g)** The number of significant intron-associated fold changes in the HEK and HeLa transfections. **h)** The EGFP and dTOM reads for each construct in HeLa, averaged across replicates. The dashed diagonal line indicates the mean GFP/dTom ratio in the intronless controls; introns that significantly increase or decrease EGFP expression relative to these controls are denoted in red and blue, respectively. **i)** The correlations in IFC across the different transfections.

**Supplementary figure 2:**
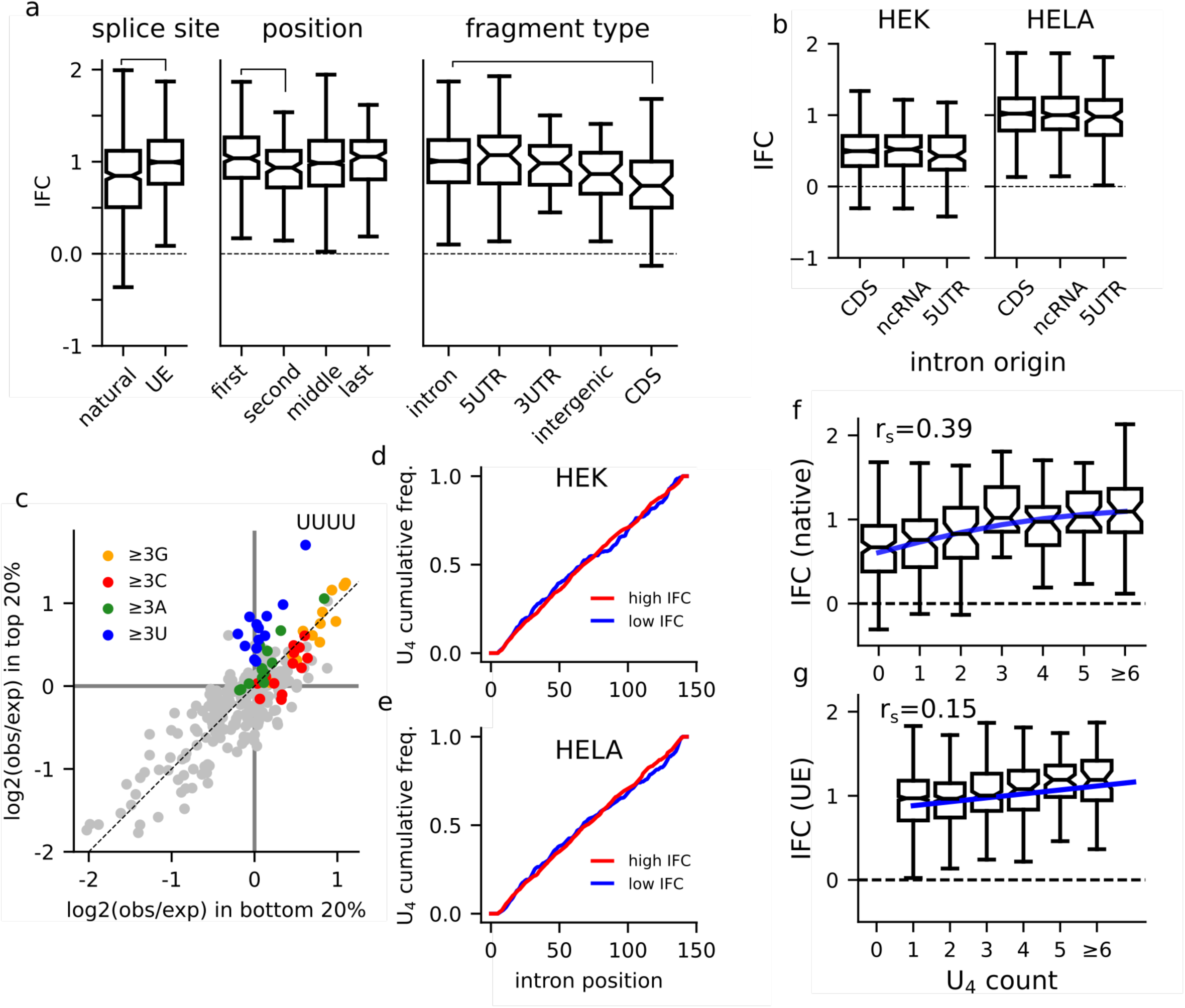
Intron features associated with IFC. **a**) Boxplots of the IFC distributions for intron bodies in the HeLa transfection broken down by native or UE splice sites; ordering within the gene; and annotated region of fragment origin. **b**) Boxplots of IFC in the HEK and HeLa transfections for natural introns originating from coding sequence, 5-UTR, or noncoding RNA. **c**) The enrichment of tetramers in introns from the top decile of IFC compared to the bottom quintile in the HeLa transfection. **d, e**) The cumulative frequency of U_4_ motif location within the intron body, for introns in the top and bottom quintiles of IFC in the HEK (**d**) and HeLa (**e**) transfections. **f, g)** The association between IFC and the number of U_4_ tetramers in the intron’s sequence for introns with native (**f**) and strong major splice sites (**g**) in the HeLa transfection; the association between U4 count and IFC is summarized with Spearman’s rank correlation (r_s_) and the trendline is a sigmoid with the steepness, midpoint, and asymptotes fit to the observations to minimize the squared error.

**Supplementary figure 3:**
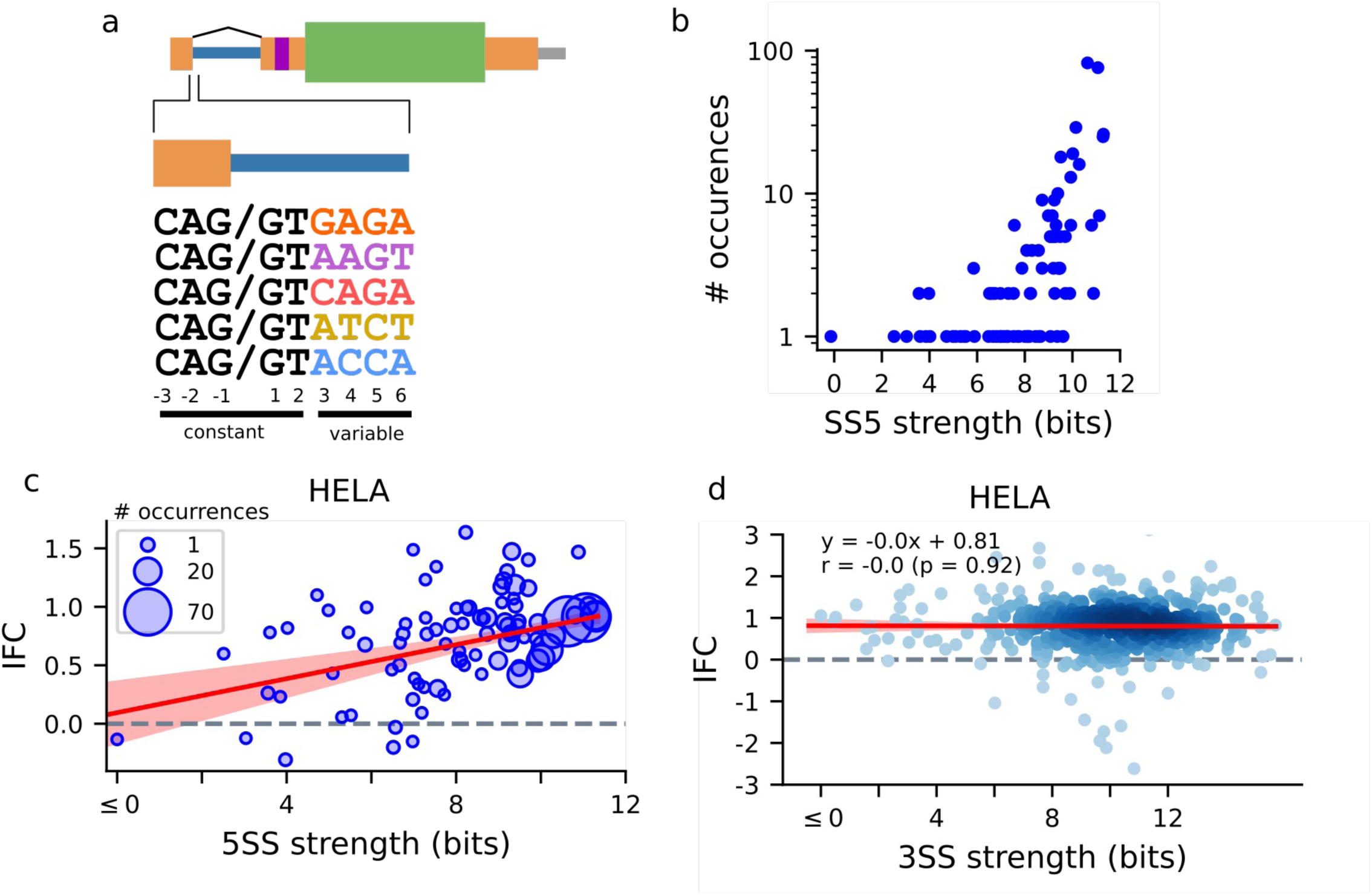
Relationship between 5SS strength, frequency and IFC in HeLa cells. **a**) Schematic of the 5SS motifs considered in the natural intron screen, highlighting the constant -3 to +2 positions and the variable +3 to +6 positions in five example motifs. **b)** The number of well-spliced natural introns in which particular 5SS motifs occur plotted against their strength. **c, d)** Top: The association between 5SS (**c**) and 3SS (**d**) strength and IME in HeLa cells, with IFC averaged across all introns with a given 5SS motif in the library; the size of the dots indicates the number of occurrences. Trend lines are linear regression fits, with 94%-intervals constructed by bootstrapping splice sites 10,000 times.

**Supplementary figure 4.**
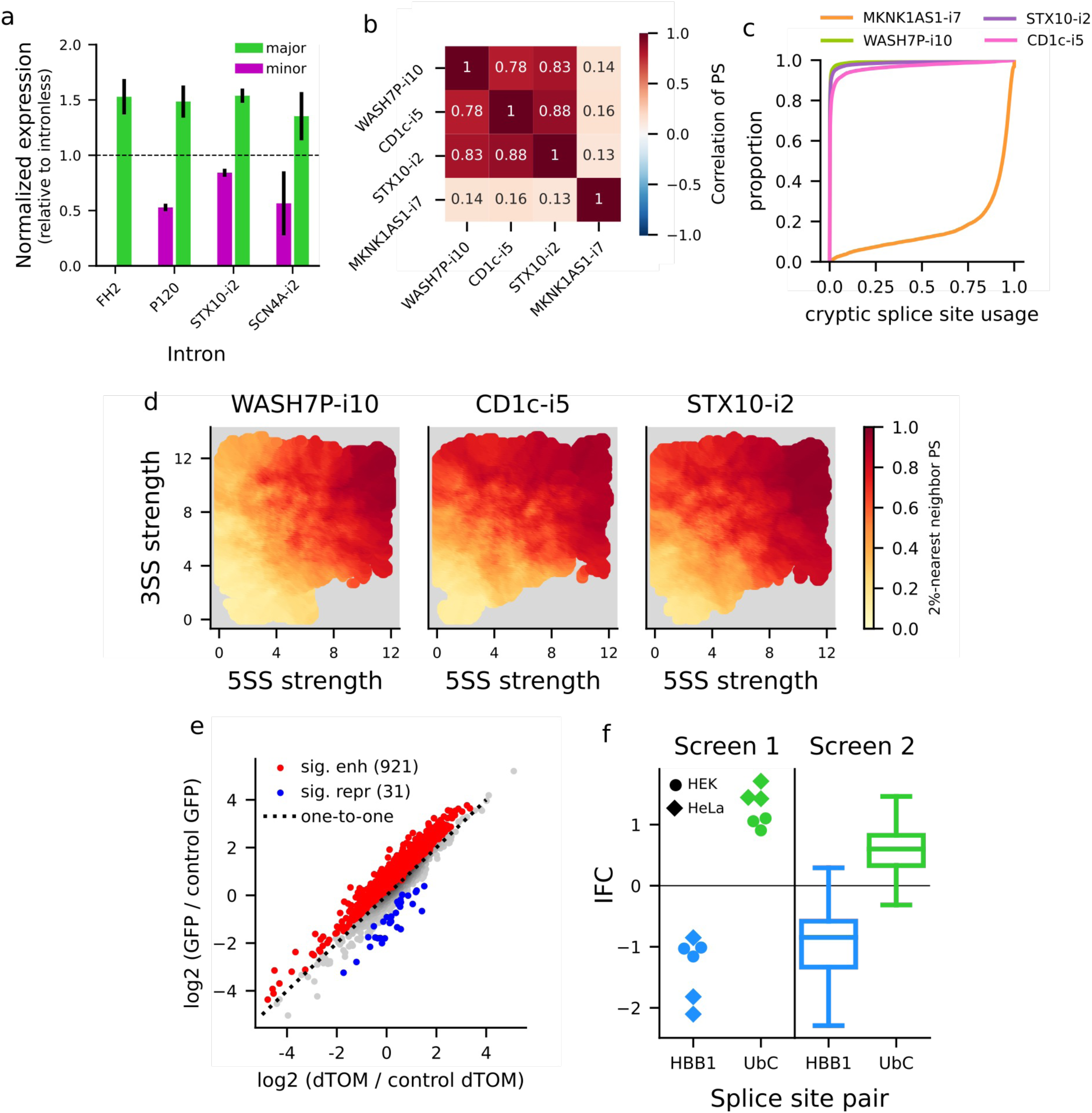
Relationship between splice site strength, splicing efficiency and IFC. **a)** qPCR measurements of the effects of intron bodies with minor or major splice sites on minigene expression, normalized to an intronless dTom control. Error bars reflect two standard errors. Dashed line indicates no difference from the intronless dTom. **b)** Correlation in PS of splice site pairs across the four intron bodies. **c)** The distribution of cryptic splice site usage rates in the four intron bodies. **d)** Relationship between the strength of splice site pairs and proportion spliced (accounting for underrepresentation of the longer, unspliced amplicons). The heatmap depicts KNN-nearest average of PS. **e)** The expression of intron-containing EGFP compared to the matched intronless dTomato, normalized to the corresponding junction-matched control reporter. Introns that significantly enhance or repress expression are colored red and blue, respectively. **f)** The effects of the major HBB1 intron 1 splice site pair on expression. Left: the IFC observed for the three replicates of HBB1-i1 in the HEK and HeLa transfections with native splice sites and with UbC splice sites. Right: The IFC associated with replacing a minor intron’s splice sites with those of HBB-i1 or UbC.

**Supplementary figure 5:**
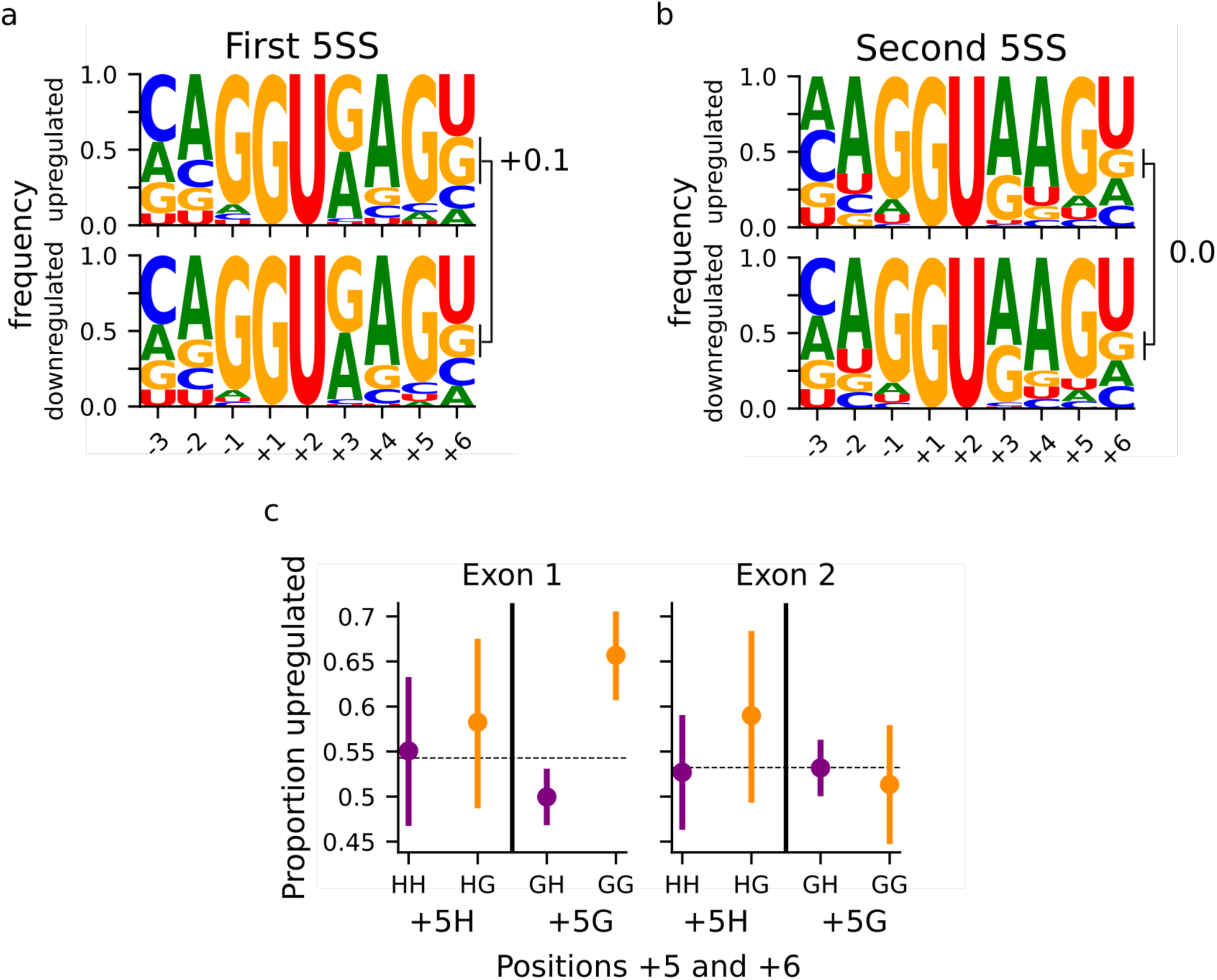
5SS features of up- and downregulated genes in U1 mutant cells. **a, b)** The position frequency matrices of the first (**a**) and second (**b**) 5SS motifs of genes reported as significantly up- and downregulated in CLL patients with the U1^3g.A>C^ mutation. c) The frequency of upregulation among significantly differential expressed genes with or without a +6G at a given 5SS motif, broken down by whether the motif also has the consensus +5G.

